# Direct neural evidence for the contrastive roles of the complementary learning systems in adult acquisition of native vocabulary

**DOI:** 10.1101/2021.04.12.439443

**Authors:** Katherine R. Gore, Anna M. Woollams, Stefanie Bruehl, Ajay D. Halai, Matthew A. Lambon Ralph

**Affiliations:** Division of Neuroscience and Experimental Psychology, School of Biological Sciences, University of Manchester, UK; St Mauritius Rehabilitation Centre, Meerbusch, & Heinrich-Heine University, Duesseldorf, Germany; Clinical and Cognitive Neurosciences, Department of Neurology, Medical Faculty, RWTH Aachen University, Aachen, Germany; MRC Cognition & Brain Sciences Unit, University of Cambridge, UK

**Keywords:** ageing, fMRI, language, semantics, vocabulary learning

## Abstract

The Complementary Learning Systems (CLS) theory provides a powerful framework for considering the acquisition, consolidation and generalisation of new knowledge. We tested this proposed neural division of labour in adults through an investigation of the consolidation and long-term retention of newly-learned native vocabulary with post-learning functional neuroimaging. Newly-learned items were compared to two conditions: (i) previously known items to highlight the similarities and differences with established vocabulary; and (ii) unknown/untrained items to provide a control for non-specific perceptual and motor-speech output. Consistent with the CLS, retrieval of newly-learned items was supported by a combination of regions associated with episodic memory (including left hippocampus) and the language-semantic areas that support established vocabulary (left inferior frontal gyrus and left anterior temporal lobe). Furthermore, there was a shifting division of labour across these two networks in line with the items’ consolidation status; faster naming was associated with more activation of language-semantic areas and lesser activation of episodic memory regions. Hippocampal activity during naming predicted more than half the variation in naming retention six months later.

## Introduction

Across the lifespan, humans need to acquire new knowledge and do so rapidly with relative ease. One lifelong learning process is vocabulary acquisition. Beyond the initial influx of new language in childhood, there are numerous words, meanings and expressions to learn throughout adulthood. Thus, individuals constantly acquire new vocabulary relating to their everyday lives, hobbies and profession. Re-establishing vocabulary is also a key target for those with language impairment (aphasia) after brain damage from injury, stroke or dementia because word-finding difficulties (anomia) are a pervasive and frustrating feature of all types of aphasia (Rohrer et al. 2008). Therefore, from both cognitive and clinical neuroscience perspectives, it is fundamentally important to understand both the cognitive and neural bases of vocabulary acquisition.

One influential theory is the Complementary Learning Systems (CLS; Marr, 1971; McClelland, McNaughton, & O’Reilly, 1995) model. This theory proposes that new knowledge is initially coded through rapidly formed, sparse representations supported by the medial temporal lobes (MTL) and hippocampus. Longer-term consolidation and evolution of generalizable representations follow from slower, interleaved learning and MTL replay to neocortical regions. Thus, over time, there is a gradual shift in the division of representational load between MTL and neocortical regions (with the rate of change depending on various factors: cf. McClelland et al., 2020). The CLS provides a potentially generalizable theoretical framework for the acquisition of many different kinds of knowledge including language acquisition (cf. Davis and Gaskell, 2009). There is, however, little direct neural evidence for this theory in long-term language learning, particularly in adults who already have large and varied vocabularies.

To date, few if any studies have explored the processes that underpin new vocabulary learning within adults’ native language (i.e., learning the meaning and name of novel items/concepts as one might do when learning about a new hobby, profession, or technology). Instead, the handful of pre-existing investigations have typically focused on second language learning. Studies have adopted different experimental designs. Some have required participants to link brand new names to pre-existing, well-established meanings (Raboyeau et al. 2004; Yang et al. 2015). Alternatively, to avoid the unfamiliar phonetic and phonological elements of second languages, researchers have used pseudowords that conform to the phonological structure of the native language (Mestres-Missé et al. 2008; Davis et al. 2009; Paulesu et al. 2009; Ozubko and Joordens 2011; Pohl et al. 2017). Of course, learning additional names for pre-existing items may generate competition between new and old words when naming. This proactive interference can skew accuracy and reaction times (Gaskell and Dumay 2003). To avoid these issues, researchers sometimes use abstract (i.e., meaningless) images alongside pseudowords. However, to understand native vocabulary acquisition and recovery of vocabulary in aphasia, investigation of meaningful real-world items with native language names would be ideal. A potentially suitable approach comes from a series of MEG and aphasiological studies that used the ‘Ancient Farming Equipment’ learning paradigm, which provides a line drawing, a novel name and a description of how the item is used (Laine and Salmelin 2010).

Existing studies of paired-associate pseudoword learning (typically assessed through recognition rather than production) suggest some important target brain regions for investigating native vocabulary. There is strong neural evidence of initial hippocampal encoding of pseudowords (Breitenstein et al. 2005; Davis et al. 2009) as well as hippocampal modulation during online pseudoword learning (Breitenstein et al. 2005). Such studies also offer neocortical regions of interest for further investigation including the left temporal lobe (Raboyeau et al. 2004; Davis et al. 2009), bilateral anterior temporal lobes (Grönholm et al. 2005) and fusiform gyrus (Breitenstein et al. 2005).

In the present study, we generated a direct evaluation of the CLS with respect to native vocabulary acquisition. Accordingly, we used fMRI to investigate the similarities and differences in neural networks and mechanisms that underlie native vocabulary learning versus well-established words. Participants were trained on novel native words for three weeks, before performing both picture naming (of previously known items, untrained/unknown items and select trained items which had been learned successfully per participant) and semantic judgement tasks in the scanner (i.e., names had to be learned sufficiently well for speech production rather than simply above-chance memory recognition). We also adopted this method and learning target as it directly mimics those found in rehabilitation of aphasic word-finding difficulties (where patients aim to re-establish meaningful, native vocabulary through multiple learning sessions, extending over several weeks). Consequently, not only does the current study provide information about native vocabulary acquisition in the healthy brain, but it may also give important clues about the neural bases of successful aphasia rehabilitation by providing a baseline for the same analysis in patients with aphasia.

We correlated reaction times (RTs) with BOLD activity from two contrasts (known > untrained and trained > untrained). These two contrasts allow testing of the main hypothesis, that differing levels of vocabulary acquisition consolidation are supported by different neural mechanisms. Items which were previously known (e.g., dragonfly, xylophone, hairdryer) are fully consolidated, whereas newly trained items (for a maximum of three weeks e.g., echidna, dilruba, binnacle) have varying levels of consolidation. Therefore, naming newly trained words versus naming previously known words may present with differing networks of neural activity, supported by the episodic or semantic system dependent on consolidation and stage of the CLS framework.

## Materials and Methods

### Participants

Twenty older, healthy native-English speakers were recruited (twelve females, age range 46-76 years, mean age 63.90, SD 8.82). All participants were right-handed, with normal or corrected-to-normal vision, no history of neurological disease, dyslexia or contraindications to MRI scanning. The Addenbrooke’s Cognitive Examination Revised (ACE-R) was used to screen for dementia, with a cut-off score of 88. Capacity for verbal learning was tested with the California Verbal Learning Test (CVLT). All participants gave informed consent before participating and the study was approved by a local National Health Service (NHS) ethics committee.

### Stimuli

There were three sets of stimuli items. All sets contained real-world items including mammals, fish, birds, tools, food, clothing and toys. Two sets included unfamiliar items with very low word frequency names. These items were drawn from the British National Corpus (BNC; Davies, 2004), a 100-million-word text corpus. One set was used for training, whilst the other remained as an untrained baseline set. The third set contained familiar items. These items were drawn from the International Picture Naming Project. Items were selected that could be named accurately (85-100%), with low word frequency and longer reaction times (>1000ms) to select less easily named items. All stimuli were below a word frequency of 100 words per 100-million and had high name agreement. In the picture naming task, fMRI stimuli were high quality, coloured photographs with a white background. In the semantic decision task, the fMRI stimuli were presented as an orthographic written name, in black text on a white background.

### Procedure

There were five stages: baseline naming assessment, word training, post-training behavioural assessment, functional imaging data collection and maintenance naming assessment (Figure 1). Participants were tested on all items before training. Stimuli sets were tailored to each participant so that all known items could be named, and all untrained and to-be-trained items could not be named prior to training. Participants undertook fMRI scanning within 2 days of finishing training. Only items which had been successfully learned, demonstrated in the post-training naming assessment, were used in the fMRI trained condition (therefore there were different stimuli sets per participant for the trained condition). To assess maintenance, participants were tested on learned items between five- and six-months post scanning, without interim training.

**Figure 1.**
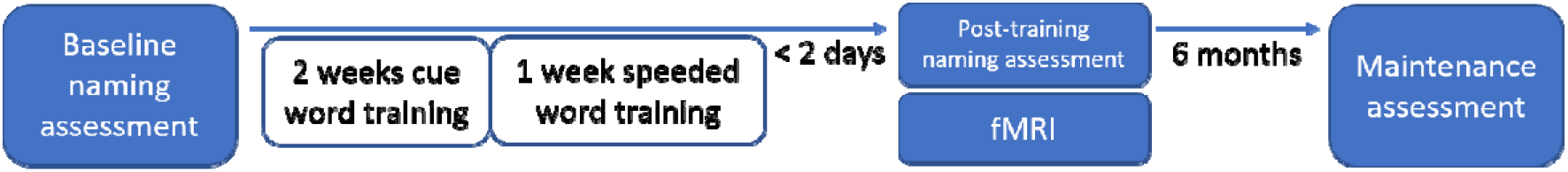
Timeline of study stages.

### Behavioural training

Participants received self-guided, at-home training on new words and the related semantic information. Training took place for up to 45 minutes a day, four days a week for three weeks. In the first two weeks, participants received cue training. In the third week, participants received speeded training.

Items were presented via an interactive PowerPoint presentation. Visual Basic for Applications was used to store cue choice, time on task and accuracy data. In weeks 1 and 2, cue training took place daily. A novel picture was shown, with the name both in orthographic and audio form. Participants were instructed to listen to the name and repeat it out loud. After all items had been repeated the cue training began. Participants were instructed only to use cues when they needed one and reminded they would be tested on the semantic information. The training was designed to allow healthy participants to choose the level of cue they thought they would need to be correct on each trial. This interactive and self-determined approach allowed the training to feel challenging, engaging and reduce boredom.

The cue training was commonly used in standard speech and language therapy (Nickels 2002; Abel et al. 2005; Pohl et al. 2017). Participants saw a picture of an item with a choice of four cues, or the option to name the item with no cues. Participants could use as many cues as they like, in any order. There were four increasing cues. First, a picture plus a written descriptive semantic cue. Second, the picture plus the first name phoneme. Third, the first and second name phoneme were cued. The fourth cue was the whole name. All cues were given both orthographically and audibly. The semantic cue was formed in the same way for each item, initially with the geographical origins, then an identifying feature, followed by a broader semantic cue. For example, an ankus was “An Indian hooked tool used to handle and train elephants.”

After each naming attempt, the whole correct word was given. Participants were asked to indicate whether they named each item correctly or not. Participants then indicated whether the item was European or not. The initial training set was 10 items. When participants were able to name 70% of the presented items with no cue, then another 10 items were added to the set, incrementally up to 50 items.

In the third week of training, the learned items were used in a novel repeated increasingly-speeded presentation (RISP; Conroy, Sotiropoulou Drosopoulou, Humphreys, Halai, & Lambon Ralph, 2018) learning environment. Participants were instructed that the computer would present an item for a short time and they needed to name the picture before a specified time limit. When participants reached a success rate of 70% at a target speed, the timing was incrementally reduced from 1.8s to 1.4s, to 1s. When participants beat the 1s target for 70% of items, the set size was increased by 10 items and the timing was reset to 1.8s.

We assessed participants’ learning using a post-training assessment of trained items in the absence of cues. Only successfully named items were used during the fMRI naming task (trained vocabulary condition; M = 45 items), creating participant-specific trained condition naming sets. The fMRI session took place on the same day as the post-training assessment.

### Neuroimaging acquisition

All scans were acquired on a 3T Phillips Achieva scanner, with a 32-channel head coil with a SENSE factor of 2.5. High resolution, whole brain, structural images were acquired including 260 slices with the following parameters: TR = 8.4ms, TE = 3.9ms, flip angle = 8 degrees, FOV = 240 × 191mm, resolution matrix = 256 × 206, voxel size = 0.9 × 1.7 × 0.9mm.

We opted to use a triple gradient echo EPI sequence in order to improve the signal-to-noise ratio (SNR), particularly in the anterior temporal lobes (ATLs) where traditionally there are issues of EPI signal dropout and distortion (Poser et al. 2006; Halai et al. 2014, 2015). All functional scans were acquired using an upward tilt up to 45 degrees from the AC-PC line to reduce ghosting artefacts from the eyes into the temporal lobes. The sequence included 31 slices covering the whole brain with TR = 2.5s, TE = 12, 30 and 48ms, flip angle = 85 degrees, FOV = 240 × 240mm, resolution matrix = 80 × 80, and voxel size = 3.75 × 3.75 × 4mm.

All stimuli were presented electronically using E-Prime 2.0 software (Psychology Software Tools). The block order was pseudo-randomised optimised for statistical power using OptSeq (http://surfer.nmr.mgh.harvard.edu/optseq/). Verbal responses were recorded using a fibre optic microphone for fMRI (FOMRI; Optoacoustics) with noise-cancelling. Participants were instructed to speak ‘like a ventriloquist’ to reduce motion artefacts.

Participants completed two tasks within during fMRI. A picture naming tasks comprised a block design with four conditions; known, trained, untrained and baseline. In the known condition, participants overtly named familiar items (e.g., umbrella). In the trained condition, participants named newly learned items (e.g, echidna). If participants could not remember an item name they responded: “Don’t Know”. If the item was novel (untrained condition), participants also responded “Don’t know”. Similarly, participants responded “Don’t know” to phase-scrambled stimuli from the other conditions as part of the baseline. The task included two trial speeds but the results did not differ across these conditions, therefore data were collapsed across this manipulation. In the standard speed condition, each 1900ms trial consisted of a fixation cross for 700ms, followed immediately by the target image in the middle of a white screen for 1200ms. With 5 items per block, each block lasted 9.5s. We also included 8 rest blocks per run which were jittered to have an average length of 9.5s. With 32 task blocks and 8 rest blocks per run, the total run time was 6 minutes and 33 seconds. In the slower condition, each trial lasted 3700ms and consisted of a fixation cross for 700ms, followed by the target image for 3000ms. Only 3 items were presented per block and each block lasted 11.1s. As before, 8 jittered rest blocks were included with an average length of 11.1s. With 32 task blocks and 8 rest blocks, the total run time was 7 minutes and 4 seconds.

The second task was required participants to make semantic decisions. This included three blocked conditions; trained, untrained and baseline. In the trained and untrained condition, participants responded “Yes” or “No” or “Don’t Know” to the semantic question “Is it European?”. In the baseline task, participants responded “Up” to an ascending alphabetical sequence “ABCD” or “Down” to a descending alphabetical sequence “DCBA”. As above, we used two trial speeds but found no differences between conditions; therefore, data were combined. In the standard speed condition, a fixation cross was displayed for 700ms, followed immediately by the target image for 1200ms (total trial = 1900ms). There were 5 trials per block each lasting 9.5s, with 6 jittered rest blocks averaging to 9.5s. The total run time was 6 minutes and 33 seconds, which included 24 task and 6 rest blocks. In the slower condition, displayed the target image for 3000ms (total trial = 3700ms). A total of 24 task blocks were used with 3 trials per block (11.1s) and 6 jittered rest blocks averaging to 11.1s (total run time = 7 minutes and 4 seconds).

### Neuroimaging pre-processing and analysis

T1 data was pre-processed using the FMRIB Software Library (FSL, version 6.0.0; http://fsl.fmrib.ox.ac.uk/fsl/fslwiki/, Woolrich et al., 2009). Brain tissue was extracted from the structural images (BET; Smith, 2002) and an initial bias-field correction was applied using FSL’s anatomy pipeline (FAST; Zhang, Brady, & Smith, 2001), excluding sub-cortical segmentation as this was performed with BET. Registration to standard space was performed in FSL with FLIRT and FNIRT (Woolrich et al. 2009; Patenaude et al. 2011) and segmentation with FAST (Zhang et al. 2001). Despiking and slice time correction was applied to the functional data in the AFNI neuroimaging suite (v19.2.10; Cox, 1996; Cox & Hyde, 1997; 3dDespike; 3dTshift). Combined normalisation, co-registration and motion correction parameter sets were applied to each functional echo in FSL. Functional data were optimally combined, taking a weighted summation of the three echoes, using an exponential T2* weighting approach (Posse et al. 1999) and regression analysis. Functional runs were also combined and denoised using multi-echo independent component analysis (ME-ICA; Kundu et al., 2013; Kundu, Inati, Evans, Luh, & Bandettini, 2012) using the tool meica.py (v3.2) in AFNI (Cox, 1996; Cox & Hyde, 1997). The denoised timeseries were normalised to standard space using FNIRT warps, then smoothed.

Statistical whole brain and region of interest analyses were performed using SPM12 (Wellcome Trust Centre for Neuroimaging) and MarsBaR. Regions of interest (ROIs) were based upon previous literature. Medial temporal lobe structures, including bilateral hippocampi are critical for episodic memory, as evidenced by hippocampal amnesia (Dickerson & Eichenbaum, 2010). However, episodic memory processes also involve the inferior parietal lobe (IPL), despite parietal lesions not resulting in episodic memory deficits (Cabeza et al. 2008). The left inferior frontal gyrus (IFG) is considered critical in speech production and semantic processes (Blank et al. 2002; Hickok and Poeppel 2007; Lazar and Mohr 2011; Price 2012). The middle temporal gyrus (MTG) is activated during semantic processing (Binder et al. 2009; Visser et al. 2012; Noonan et al. 2013; Jackson 2021), and focal damage is associated with semantic deficits (Dronkers et al. 2004). The specific co-ordinates for these ROIs were derived by conducting a Neurosynth (Yarkoni et al. 2011) fMRI meta-analysis using two search terms: ‘episodic memory’ (bilateral hippocampi; MNI: −28 −14 −15, 29 −14 −15 and left IPL MNI: −47 −64 34), and ‘language’ (left IFG MNI: −46 28 10 and left MTG MNI: −52 −42 0).

Furthermore, we included a left ventral anterior temporal lobe (vATL) ROI (MNI: −36 −15 −30) taken from a key reference (Binney, Embleton, Jefferies, Parker, & Lambon Ralph, 2010). The vATL is often missed in fMRI studies using typical echo times of >30ms at 3T due to signal dropout. However, there is clear evidence from the neuropsychology literature and semantic dementia patients that the vATLs are important for semantic cognition patients (Rogers et al. 2004; Patterson et al. 2007; Lambon Ralph 2014; Lambon Ralph et al. 2016). Indeed, there is growing evidence that fMRI protocols optimised for signal detection in areas of magnetic susceptibility can identify vATL areas during semantic processing (Devlin et al. 2000; Halai et al. 2014, 2015; Jackson et al. 2016; Rice et al. 2018).

Reaction times (RTs) for ROI analyses were calculated from onset of stimulus and were z-scored to account for any variance due to time on task. RTs were z-scored by condition to enable analysis of within-condition RT variance.

## Results

### Behavioural Data

Participants spent a mean of 4.3 hours training (SD = 0.8) over an average of 12 sessions. Participants successfully learned the novel vocabulary with an average gain of 81% (SD = 10.73) outside the scanner. Inside the scanner, in the trained condition, participants were presented with only items they had successfully learned during training, ascertained by a post-training behavioural picture naming task. Participants had an average of 88% (SD = 12.0) accuracy on these participant-specific trained items in scanner and an average reaction time of 1054ms (SD = 203.5) in scanner. Participants also successfully learned semantic information about these successfully trained novel items, with an average gain of 83% (SD = 12.92). This level of variation in semantic knowledge of the new items (“Is it European?”) demonstrates there was a continuum of semantic consolidation between participants in the trained items. As would be expected, naming accuracy for already-known (pre- and post-training) items during scanning was high (M = 98%, SD = 2.2), with a mean reaction time of 1020ms (SD = 148.5). The naming latency for the newly learned and previously known items were not significantly different (t(19) = −.396, p = .696), indicating effectiveness of the training.

To explore the effect of semantic knowledge on word learning, correlations were performed between accuracy in the semantic judgement task (“Is it European?”), naming accuracy and subsequent maintenance of naming accuracy. There was a significant positive correlation between naming and semantic accuracy, with age at scan added as a controlled variable (r(20) = .912, p = .000). This correlation suggests that acquisition of semantic knowledge and names is bound together and, potentially, that semantics facilitates word learning. Additionally, there was a significant correlation between learning of the semantic cues and overall maintenance of learned, trained items (*r*(20) = .66, p = .002), suggesting that a deeper encoding of knowledge is key to long-term word learning.

### Whole brain results

The results of the whole brain analyses are reported in Table 1, where three contrasts were created: 1) trained > untrained, 2) known > untrained and 3) trained > known. There were no significant clusters of activation for the opposing contrasts: untrained > trained, untrained > known and known > trained. There was a similar pattern of activation between the contrasts, where large bilateral language areas where identified. There was, however, greater and more extensive activation for the trained condition, including the hippocampus in both the trained > untrained and trained > known contrasts (Figure 2). The increased activity may be due to increased task demands for the trained condition, as evidenced by increased RTs, but it is of note that the episodic-hippocampal activity was for trained items only, in both the trained > untrained and trained > known contrasts.

**Table 1.**
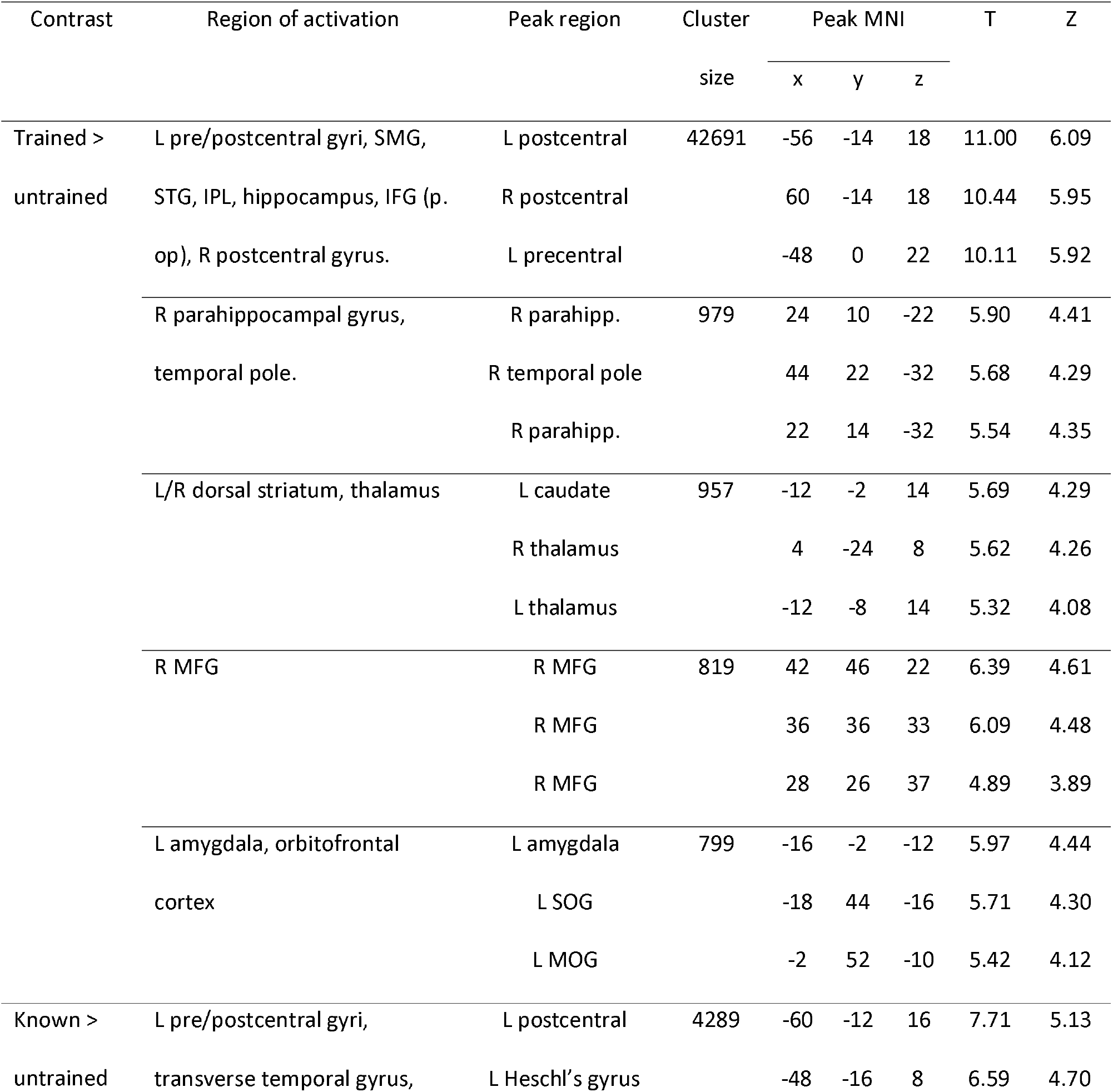
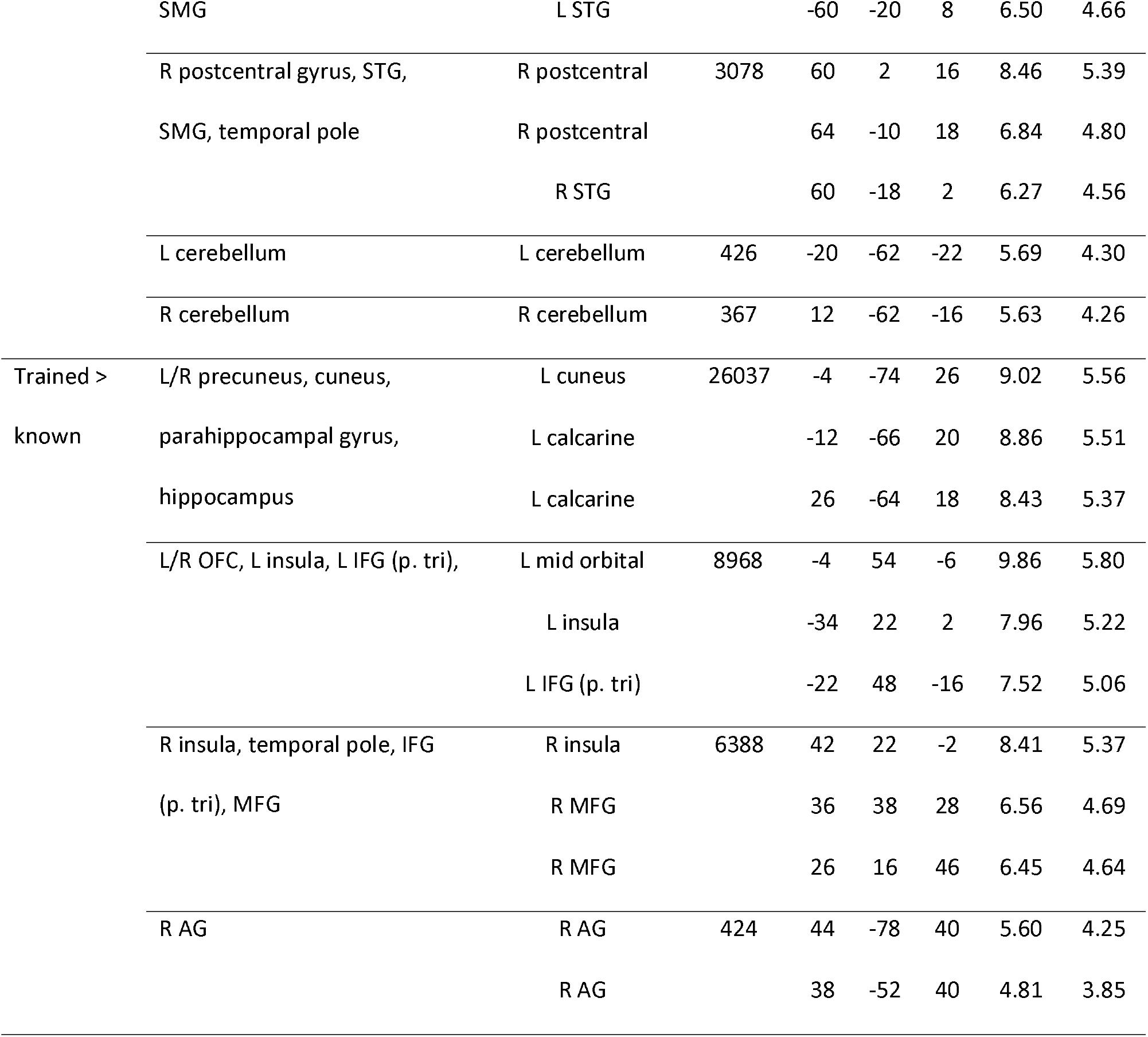
Clusters significant at p< 0.001 voxel height and p < 0.05 FWE cluster correction for picture naming trained, known and untrained items. Up to 3 strongest peaks listed per cluster, peak MNI = x, y, z. L; left, R; right, SMG; supramarginal gyrus, STG; superior temporal gyrus, IPL; inferior parietal lobe, IFG op; inferior frontal gyrus pars opercularis, IFG tri; inferior frontal gyrus pars triangularis, OFC; orbitofrontal cortex, MFG; middle frontal gyrus, AG; angular gyrus, parahipp; parahippocampal gyrus, SOG; superior orbital gyrus, MOG; middle orbital gyrus.

**Figure 2.**
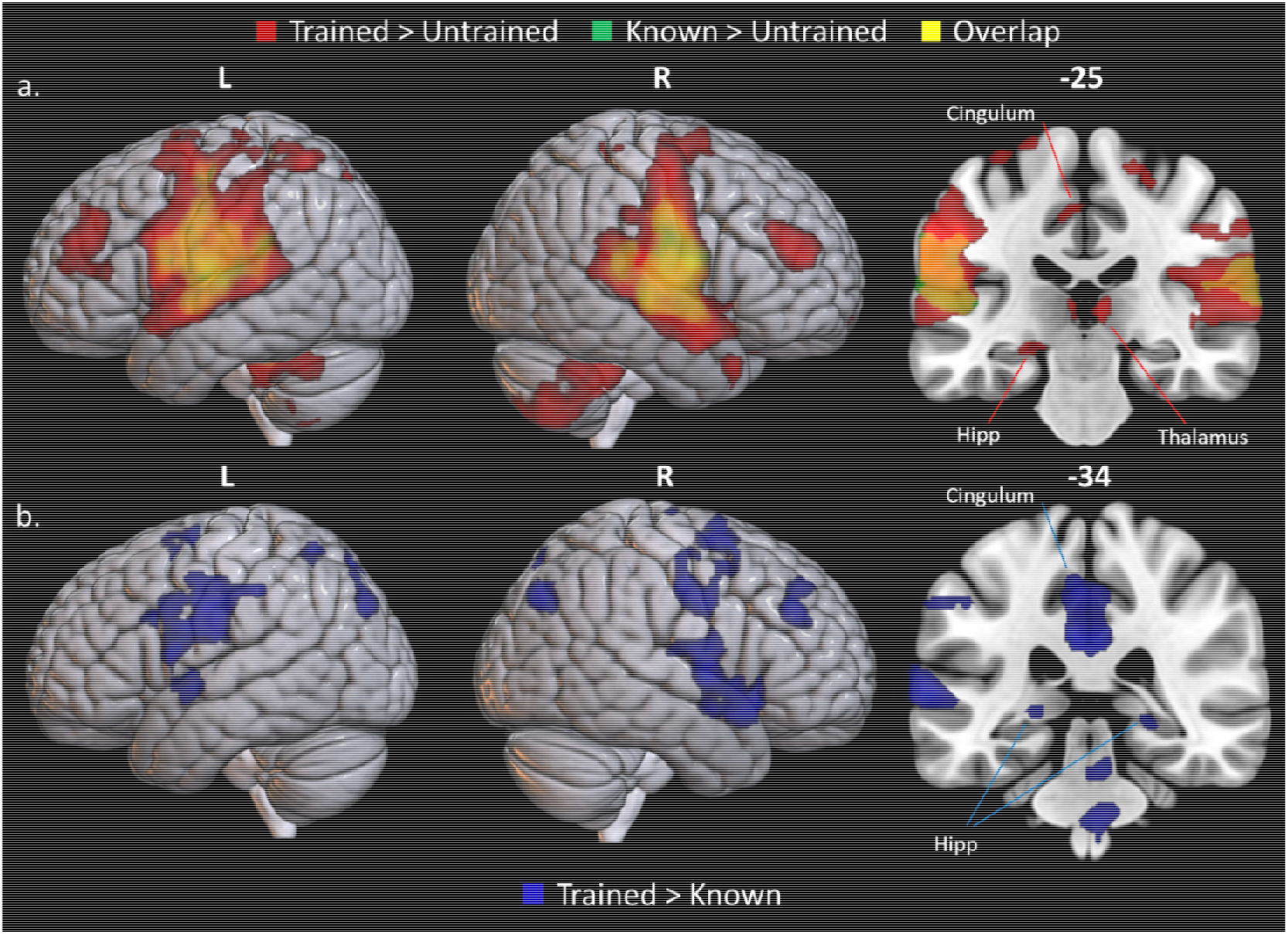
Whole brain BOLD activation of picture naming; a) trained minus untrained items (red) and known minus untrained items (green); yellow = overlap. b) trained minus known items (blue). Images thresholded at *p* < 0.001 voxel height, FWE-cluster corrected *p* < 0.05. L; left, R; right, Hipp; hippocampus, Thal; thalamus.

We did not find any significant clusters for the trained-untrained contrast in the semantic task; however, a significant cluster was identified for the untrained-trained contrast in the right anterior temporal lobe (rATL; peak MNI: 52, 4, −15) extending to the right superior temporal gyrus (STG; peak MNI: 56, 2, −12).

### ROI analysis

To explore a core hypothesis arising from the CLS theory (a division of labour between MTL vs. cortical regions), behavioural data were correlated with activity in a priori ROIs related to episodic memory (bilateral hippocampi and left inferior parietal lobe; IPL) and semantic memory (left inferior frontal gyrus; IFG, left middle temporal gyrus; MTG and left anterior temporal lobe; ATL; Figure 3b). There were no significant correlations between semantic behavioural performance and *a priori* ROIs.

**Figure 3.**
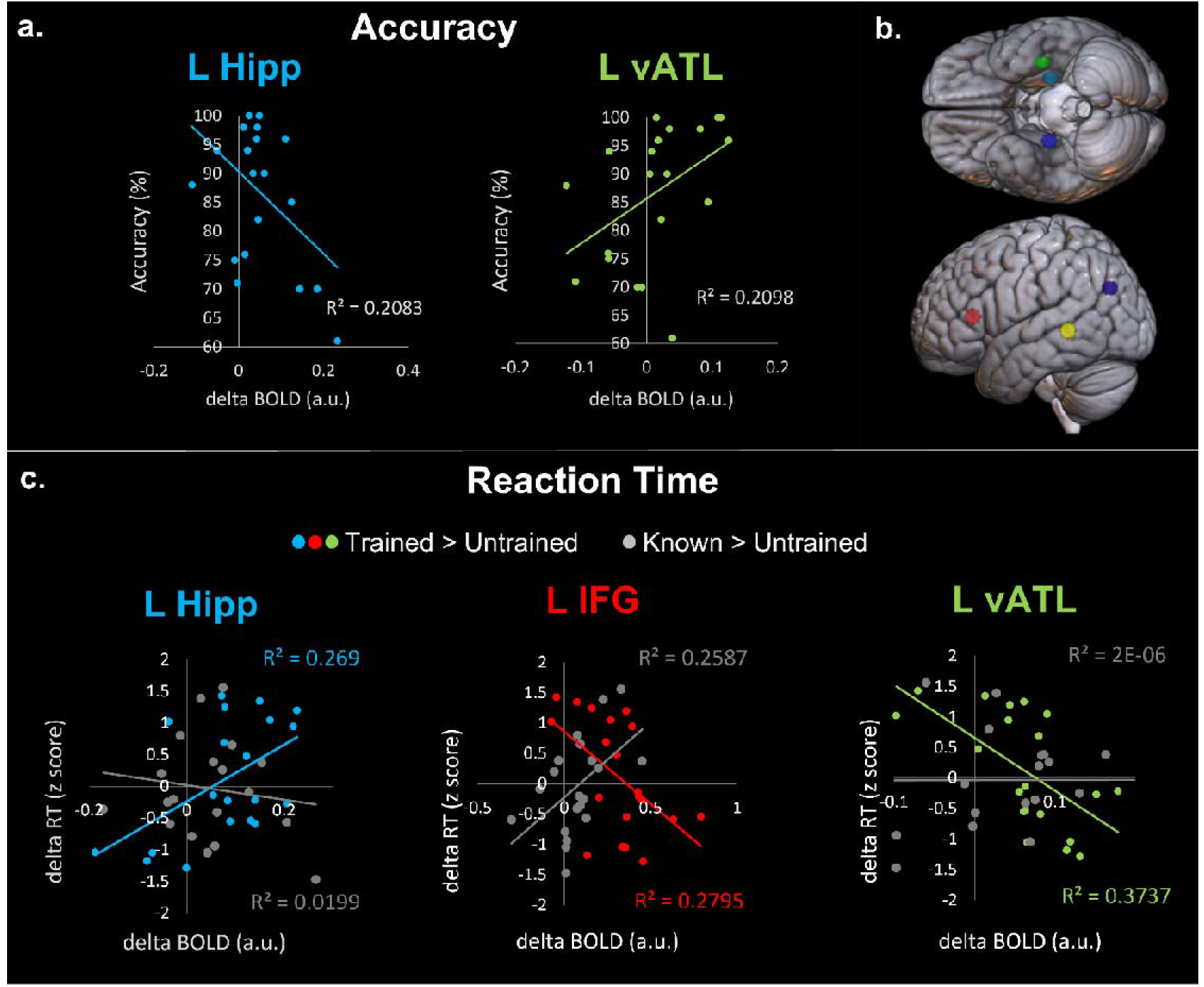
a) Significant correlations of post-training percentage accuracy of trained items versus average BOLD for trained > untrained contrast. b) Spherical 6mm regions of interest: right hippocampus (navy; MNI: 28 −14 −15) left hippocampus (cyan; MNI: −28 −14 −15), left inferior parietal lobe (purple; IPL, MNI: −47 −64 3), left inferior frontal gyrus (red; IFG, MNI: −46 28 10), left anterior temporal lobe (green; vATL, MNI: −36 −15 −30), left middle temporal gyrus (yellow; MTG, MNI: −52 −42 0). c) Significant correlations between contrast estimates (pink; trained > untrained, purple; known > untrained) and normalised in-scanner RT per participant per condition.

In the initial exploratory analysis, for the trained > untrained contrast, we found a positive correlation between the left hippocampus and longer RTs (*r* = .519, p = .019; Figure 3c). Conversely, we observed inverse correlations in semantic areas located in IFG (IFG; *r* = −.528, p = .017; Figure 3c) and ATL, (*r* = −.611, p = .004; Figure 3c), where greater activation was related to quicker performance, suggesting they had deeper consolidation in the corresponding neocortical regions. There were no further correlations between trained > untrained BOLD and naming RTs. In the known > untrained contrast, there was a significant correlation between RT and left IFG BOLD activity (*r* = .509, p = .022; Figure 3c). There were no further significant correlations between known > untrained BOLD activity and RT, including the left hippocampus (*r* = −.106, p = .656) and left ATL (*r* = .001, p = .995; Figure 2c).

The key test of the CLS hypothesis is whether the trained > untrained behavioural correlations were significantly different from the known > untrained correlations, indicating differing neural networks for naming fully consolidated known items, versus less consolidated newly trained items (for a maximum of three weeks). The correlations of hippocampal activity in the trained > untrained contrast and RT, versus hippocampal activity in the known > untrained contrast and RT were significantly different using Fisher’s r-to-z transformation (z = 1.846, p = .032, adjusted p = .032), in addition to the left ATL correlations (*z* = −2.25, p = 0.012, adjusted p = 0.018) and the left IFG correlations (*z* = −3.348, p = .001, adjusted p = 0.001), Benjamini-Hochberg adjusted p values for multiple comparisons (Benjamini and Hochberg 1995), *p* = 0.05.

We also correlated in-scanner accuracy with BOLD activity for the trained > untrained contrast in each ROI to give a measure of overall word acquisition and learning robustness. In the left hippocampus, individuals with greater activity showed poorer learning (*r* = −.456, p = .043; Figure 3a). Conversely, greater activity in the left ATL related to better accuracy (*r* = .450, p = .046; Figure 3a). Previously known items could be correctly named on three separate behavioural testing occasions, therefore, there was high (M = 98%) accuracy on these items in the scanner, which does not provide variation for correlation with BOLD activity and therefore negates the ability to test the key hypotheses. These two correlations were significantly different to each other however, using Fisher’s r-to-z transformation (*z* = −2.85, p = .004). All other correlations for trained > untrained accuracy, and known > untrained accuracy, with the a priori ROIs were not significant.

### Maintenance data

Participants were re-tested on learned items five to six months post scanning, without interim training. Maintenance varied across participants, but overall participants named on average 73.9% (SD = 27.43) of learned words. To explore whole-brain correlates, percentage drop-off in naming performance over the maintenance period was added as a covariate of interest to the trained > untrained contrast. We identified a cluster in the right hemispheric dorsolateral prefrontal cortex (rDLPFC, peak MNI: 38 8 46) corresponding to better maintenance (Figure 4a). There was no significant difference in the opposing direction or for the known-untrained contrast for either group.

**Figure 4.**
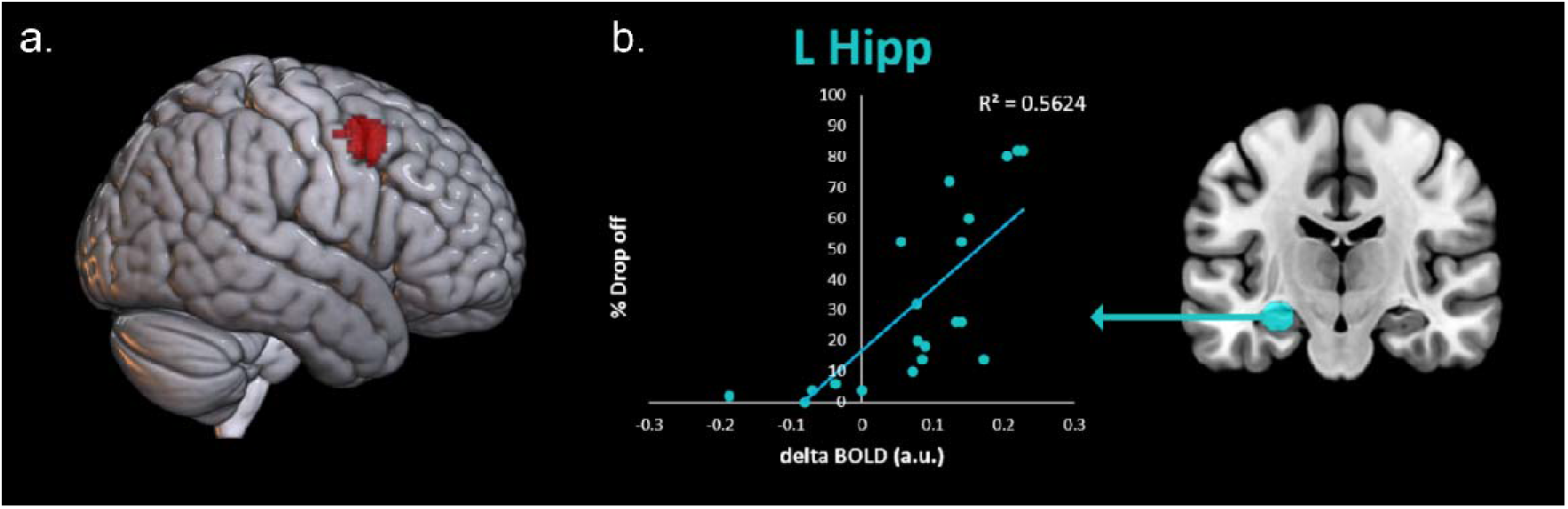
Correlations between maintenance and brain data. a) Percentage maintenance drop off as a covariate of interest in the trained-untrained contrast. Image thresholded at p < 0.001 voxel height and p < 0.05 FWE-cluster correction. b) Significant correlation between left hippocampal activity and percentage drop off.

To explore the predictions from the CLS framework, we obtained a correlation between maintenance and the *a priori* ROIs during naming of trained words (Figure 4b). There was a significant positive correlation between left hippocampal activation and percentage drop off (r(20) = .605, p = .005), which suggests that individuals who were more reliant on hippocampal structures after learning, were less likely to retain the newly-learned vocabulary after a delay. There were no other significant correlations for trained > untrained or known > untrained contrasts.

## Discussion

Vocabulary acquisition is a lifelong process for everyday life (e.g., ‘coronavirus’), hobbies (e.g., ‘thermocline’) and careers (e.g., ‘temporoparietal’). Reviving vocabulary is also key for individuals with language impairment after brain injury, stroke or dementia. This study evaluated the Complementary Learning Systems (CLS) framework (McClelland et al., 1995; McClelland et al., 2020) for the acquisition of novel real-world vocabulary in adulthood. At one time-point post-learning, a continuum of consolidation is demonstrated with responding to completely unknown and untrained words, naming successfully trained words with varying levels of semantic knowledge, and naming previously known, well consolidated items. The results indicated that: a) new learning activates a combination of the typical language-semantic network, plus the hippocampal-episodic memory network; b) activity in these network regions show opposing correlations depending on how effective learning occurred; and c) maintenance was related to consolidation of information that pertained to language networks and semantic knowledge.

### Complementary learning systems

The idea of multiple memory systems is a longstanding concept since William James argued in 1890 for “primary” (short-term) and “secondary” (long-term) memory. Multiple systems have been discussed in the context of dissociative patient studies; implicit versus explicit (Graf and Schacter 1985), declarative versus nondeclarative (Squire and Zola 1996), semantic versus episodic (Tulving 1972) and familiarity versus recollection (Mandler 1980). Exploring these disassociations gives insight into the neural structure and function of memory. However, viewing these systems as interactive and cooperating brain systems instead, allows for the ability to learn new items without catastrophic interference (McCloskey and Cohen 1989). The learning results described in this study fit perfectly with the CLS model. The CLS framework proposes a two-stage episodic-semantic account of learning; initial rapid hippocampal storage of new memories, followed typically by slower interleaved consolidation of new information alongside existing knowledge in the neocortex (McClelland, 2013; McClelland et al., 2020).

In this study, we observed hippocampal activity being related to learning new words as observed in previous studies (Breitenstein et al. 2005). In the CLS model, sparse representations in the hippocampus allow for quick learning without catastrophic interference of existing knowledge. Therefore, we predicted the role of the hippocampus should decline as learned information is consolidated into long-term neocortical representations. We indeed established that accuracy and latency immediately after learning, and long-term retention were related to the amount of hippocampal activation.

The neocortical areas activated by naming of newly learned items were typical of areas identified during speech production (Blank et al. 2002; Price 2012). We also identified two cortical regions associated with proficiency of naming learned items – the left anterior temporal lobe (ATL) and left inferior frontal gyrus (IFG). These areas are typically associated with semantic and language processing. The ventrolateral anterior temporal lobe is considered to be a trans-modal hub critical to semantic representation (Lambon Ralph et al., 2016). This proposal has strong, convergent support from multiple sources including semantic dementia patients (Acosta-Cabronero et al., 2011; Jefferies & Lambon Ralph, 2006; Patterson, Nestor, & Rogers, 2007; Warrington, 1975), fMRI (Binney, Embleton, Jefferies, Parker, & Lambon Ralph, 2010; Visser, Jefferies, Embleton, & Lambon Ralph, 2012), transcranial magnetic stimulation (Pobric et al. 2007, 2010), surface cortical electrode studies (Shimotake et al. 2015) and computational modelling (Rogers et al. 2004; Chen et al. 2017; Hoffman et al. 2018; Jackson et al. 2021). Sub regions of the LATL have been associated with picture naming and speech production specifically (Sanjuán et al. 2015). The IFG has been linked to speech production, amongst other processes, since Broca (1861) reported a patient with loss of articulation after destruction of the IFG and surrounding cortex. Despite debate as to the exact role of subregions of the IFG in speech production (Flinker et al. 2015) and semantic control (Whitney et al. 2012; Jefferies 2013; Noonan et al. 2013; Jackson 2021), the IFG is widely recognised to be important for articulation (Blank et al. 2002; Hickok and Poeppel 2007; Lazar and Mohr 2011; Price 2012).

Interestingly, we found differences between how the ATL and IFG related to accuracy and/or efficiency compared to the hippocampus. For example, faster responses and higher accuracy was related to greater ATL and IFG activity. In contrast, lower hippocampal activity was indicative of faster responses and/or higher accuracy. This provides supporting evidence for the second stage of the CLS model, where increased neocortical activity is thought to support consolidation of knowledge (indicated by improved accuracy and efficiency). We did not identify a correlation between the IFG and accuracy; but this could be due to a lack of power. It is possible that by using a high-level baseline of responding to untrained items results in multiple elements of the speech production network to be engaged. This may include activity within the left IFG which would lead to a relatively small effect size. The correlation between IFG activity and known RT may be due to less neural effort due to familiarity in terms of previous articulation, (Blank et al. 2002; Price 2010, 2012) or semantic control requirements (Jackson, 2021; Jefferies, 2013; Lambon Ralph, Jefferies, Patterson, & Rogers, 2016) for the more easily accessible known items. The known stimuli were chosen for lower word frequency, however, different words will be more or less common between individuals dependent on experience and exposure which may result in differential neural effort.

It has previously been hypothesised that the CLS could apply in other domains (Davis and Gaskell 2009) and there are demonstrations in short-term pseudoword recognition (Cornelissen et al. 2004; Breitenstein et al. 2005; Mestres-Missé et al. 2007; Davis et al. 2009). Our findings complement and significantly extend these intra-learning investigations by exploring learning after full consolidation and maintenance of the new vocabulary. With three weeks training, the participants were able to name the items without cueing and make semantic decisions (i.e., more than exhibit above-chance recognition performance). Taking this body of literature together, they clearly demonstrate that the hippocampal system is critical for new learning of artificial and native vocabulary learning, and that long-term consolidation reflect the gradual shift to long-term cortical representation and processing as predicted by the CLS model.

The speed of consolidation and reliance on the hippocampal-episodic network is now understood to be dependent on the strength of relationship between pre-existing knowledge and information to be learned (Kumaran et al., 2016; McClelland, 2013; McClelland et al., 2020). In this study, participants learned entirely new information (items, semantics and names). The item names are arbitrarily related to the object and their associated meaning; thus, this new knowledge is not systemically related to any pre-existing information. Therefore, the results obtained were as expected – it takes time to consolidate item names and, even after 2-3 weeks of learning, individuals remain reliant on a mixture of the hippocampal-episodic and semantic systems, rather than entirely on the cortical language-semantic system.

### Translational potential

This study is also potentially informative for aphasia therapy. The neural bases of successful speech and language therapy have been rarely explored, and those studies that have done so have yielded varying results (Abel et al. 2015; Nardo et al. 2017; Woodhead et al. 2017). The methods adopted in this study were deliberately designed to mimic those used to treat word-finding difficulties (where patients aim to re-establish meaningful, native vocabulary through multiple learning sessions and vanishing phonemic cues (Abel et al. 2005), over several weeks (Dignam et al. 2016). By using the same paradigm, future studies can explore whether the neural correlates of word learning/re-learning in aphasia follows the same framework. The current results would seem to imply that therapy success will depend on: (i) the extent of damage to specific critical regions involved in the CLS framework; and (ii) damage to connectivity from the hippocampus to critical language regions. Furthermore, the majority of patients (especially those with middle cerebral artery stroke) tend to have intact hippocampus, which may be linked to the reason why patients experience initial success in learning but long-term learning and maintenance (the goal of therapy) will relate to how well the therapy can induce relearning/stabilisation of neocortical representations. If these mechanisms hold in stroke aphasia, it could have important implications for intensity and dose of speech and language therapy provision.

### Conclusions

The results of the study provide a framework for word learning in the healthy older brain. We provide direct evidence of both aspects of the CLS model in long-term native word acquisition. Firstly, initial sparse hippocampal encoding with slower, less accurate and less maintenance of naming associated with episodic network activation. Secondly, consolidation to neocortical regions with rapid, more accurate and maintained naming associated with language networks. Critically, these associations are only found in new learning, not during naming of known items.

## Acknowledgements

This work was supported by a Medical Research Council National Productivity Investment Fund PhD studentship, and grants from The Rosetrees Trust (no. A1699), the European Research Council (GAP: 670428 - BRAIN2MIND_NEUROCOMP) and a Medical Research Council programme grant (MR/R023883/1) and intra-mural funding (MC_UU_00005/18). To whom correspondence may be addressed.

